# Correction of a recurrent pathogenic variant in methylmalonic acidemia using adenine base editing

**DOI:** 10.64898/2026.03.12.711365

**Authors:** Elena M. Kahn, Hooda Said, Ping Qu, Mohamad-Gabriel Alameh, Xiao Wang, Kiran Musunuru, Rebecca C. Ahrens-Nicklas

## Abstract

Methylmalonic acidemia (MMA) is a recessive genetic disease caused by variants in the *MMUT* (mitochondrial enzyme methylmalonyl–CoA mutase) gene or by defects in transport or metabolism of MMUT cofactor (5’ deoxyadenosylcobalamin), including variants in the *MMAB* gene. For the most recurrent pathogenic *MMAB* variant, c.556C>T (R186W), we identified a corrective editing strategy using adenine base editing. Deploying an adenine base editor mRNA and optimized hybrid guide RNA with lipid nanoparticles, we observed efficient *in vitro* corrective editing of the variant to wild-type, with minimized bystander editing and off-target editing in hepatocytes. These observations lay the groundwork for a gene editing therapy for patients with MMA resulting from at least one copy of the *MMAB* c.556C>T (R186W) variant, as well as a platform of similar therapies for patients with MMA caused by other variants amenable to adenine base editing.

## Introduction

Hepatic inborn errors of metabolism (IEMs) are individually rare but collectively affect 1:1000– 1:2500 births.^1,2^ Most arise from autosomal recessive loss-of function variants in genes encoding key enzymes in hepatic biochemical pathways. Loss of enzyme activity results in accumulation of upstream toxic metabolites and/or insufficient production of downstream products. In many cases, abnormal liver biochemistry induces secondary organ dysfunction, especially in the brain. Each disorder has a distinct molecular etiology, with more than 140 hepatic IEMs cataloged to date. However, many hepatic IEMs, including methylmalonic acidemias (MMA),^3^ share cardinal features that make them ideal candidates for a platform-based gene editing approach including: (1) the molecular etiology (i.e., editing target) is unambiguous; (2) accumulated metabolites are well-established disease and therapeutic biomarkers; (3) studies demonstrate the clinical benefit of liver correction (an organ that is accessible with current delivery technologies, particularly lipid nanoparticles [LNPs]); (4) restoring 10-20% of hepatic enzyme activity often corrects disease phenotypes; and (5) most patients in the U.S. are identified as neonates through universal newborn screening. Indeed, we recently reported the use of LNP-mediated, liver-directed adenine base editing to treat an infant diagnosed with neonatal-onset carbamoyl phosphate synthetase 1 (CPS1) deficiency, resulting in improved biomarkers and increased tolerance of protein in the diet.^4^

The overall incidence of MMAs varies globally, but it is estimated to be approximately 1:100,000-200,000 in the U.S. and Europe.^3^ Patients with MMA cobalamin B type (MMAB disease, MIM251110), who represent a small subset of the total MMA population, harbor biallelic pathogenic variants in the *MMAB* gene. There is a recurrent severe, infantile-onset variant, *MMAB* c.556C>T (p.Arg186Trp or R186W), that accounts for a large proportion of alleles in reports on patients with MMAB disease.^5,6^ This variant is frequently found in the homozygous state and is associated with a complete loss of protein as assessed by Western blot analysis.^7^

Humans ingest protein to support growth and the synthesis of various key macromolecules. Once ingested, the propiogenic amino acids (valine, isoleucine, methionine and threonine) are broken down by the vitamin B12-dependent enzyme, methylmalonyl CoA mutase. Patients with MMAB disease cannot fully process vitamin B12 (i.e., cobalamin) and therefore cannot activate the mutase enzyme.^3^ As such, they experience frequent episodes of life-threatening metabolic decompensation, characterized by severe acidosis due to methylmalonic acid accumulation.^8^ Patients can also develop profound hypoglycemia and hyperammonemia, causing severe neurologic damage.

The standard of care focuses on limiting intake of propiogenic amino acids and preventing catabolism through dietary measures and preventing acidosis through chronic administration of high-dose vitamin B12 and bicarbonate. Given that these measures are poorly effective, liver transplantation has become a therapeutic option for severely affected patients at many institutions.^9,10^ While liver transplantation does not fully correct all features of the disease, it greatly reduces the risk of metabolic decompensation, improves outcomes, and prolongs survival.^10^ However, transplantation is often delayed by donor availability and the need for a patient to grow to an appropriate size for transplant,^11^ and transplantation carries both acute and chronic health risks.

Given the substantial unmet medical need experienced by patients with MMAB disease, we sought to develop an adenine base editing therapy that could provide durable treatment for a subset of patients, specifically those with the c.556C>T (R186W) variant in at least one copy of the *MMAB* gene. The expected effect of reverting the variant to wild-type would be to restore functionality to the MMAB protein, durably reducing blood methylmalonic acid levels and addressing other laboratory abnormalities in patients.

## Results

To identify a base editing solution for the *MMAB* R186W variant, we adopted our previously reported strategy^4^ and used a lentiviral vector to transduce human HuH-7 hepatoma cells with a 101-bp *MMAB* genomic sequence spanning the variant, along with 101-bp genomic sequences spanning the *PAH* P281L and R408W variants. The *PAH* variant sequences, which are recurrent pathogenic variants causing PKU, serve as reference controls due to their having validated base editing solutions that we have previously shown to be efficacious in correcting the variants *in vitro* in HuH-7 cells and *in vivo* in humanized mouse models of PKU with either of the variants.^12,13^ Using the lentivirus-transduced R186W HuH-7 cells, we screened a variety of adenine base editors (ABEs) in combination with individual candidate guide RNAs (gRNAs) in plasmid transfection experiments. Seven guide RNAs (designated gRNA3 through gRNA9) tiling the site of the R186W variant, such that the variant adenine base ranges from positions 3 through 9 of the protospacer sequence (**Figure 1**) that span the ABE editing window, were tested with ABEs compatible with the protospacer-adjacent motifs (PAMs) associated with each of the protospacer sequences.^14^ ABEs with three different deaminase domains were used: ABE8.8, with the narrowest editing window, ABE8.20, with an intermediate editing window, and ABE8e, with the broadest window.^15,16^ We found that the combination of SpG-ABE8e and gRNA8 had the highest corrective editing efficiency for the R186W variant while minimizing unwanted bystander editing of nearby adenines (**Figure 2A**, red arrow).

**Figure 1.**
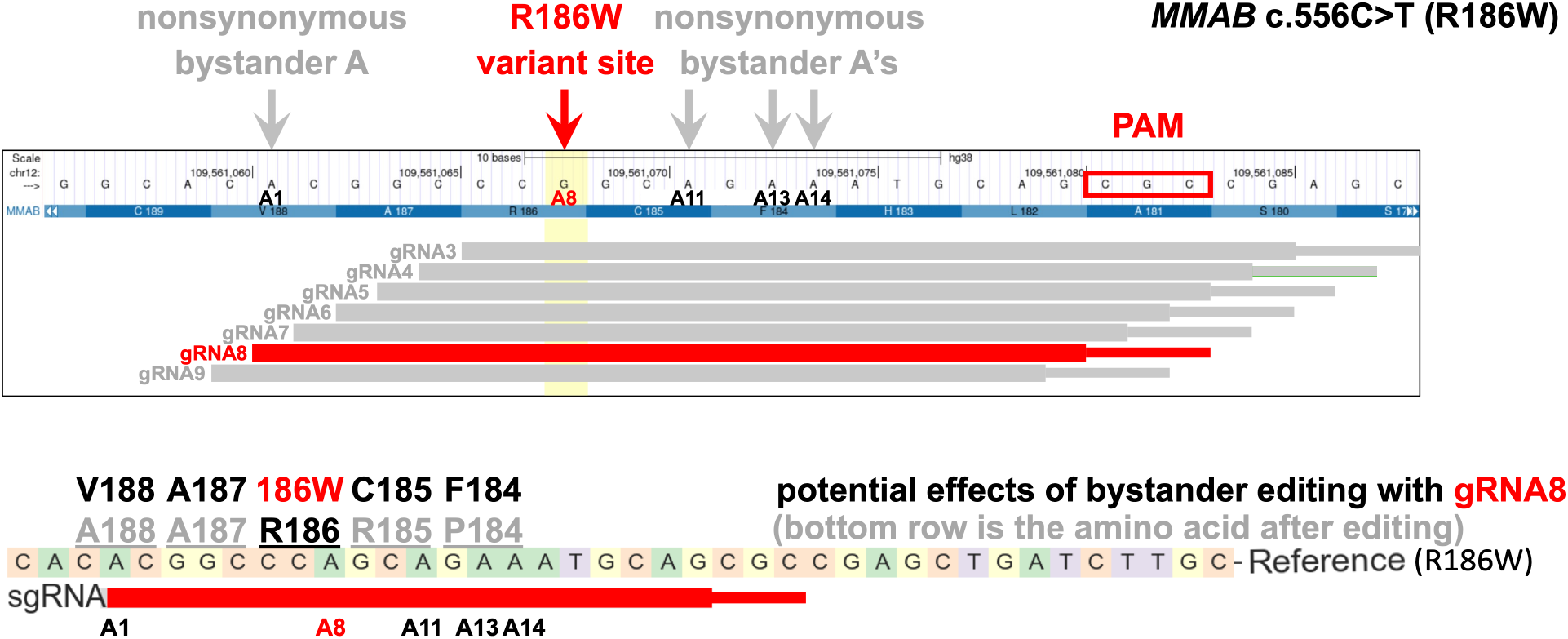
Schematic of the genomic site of the *MMAB* c.556C>T (R186W) variant. Adapted from the UCSC Genome Browser (GRCh38/hg38). The red arrow and the vertical yellow bar indicate the position of the G altered to A (A8, in red) by the variant on the antisense strand. The grey arrows indicate the sites of potential bystander editing. The horizontal bars indicate protospacer (thick) and PAM (thin) sequences targeted by the gRNA3 through gRNA9 guide RNAs.

**Figure 2.**
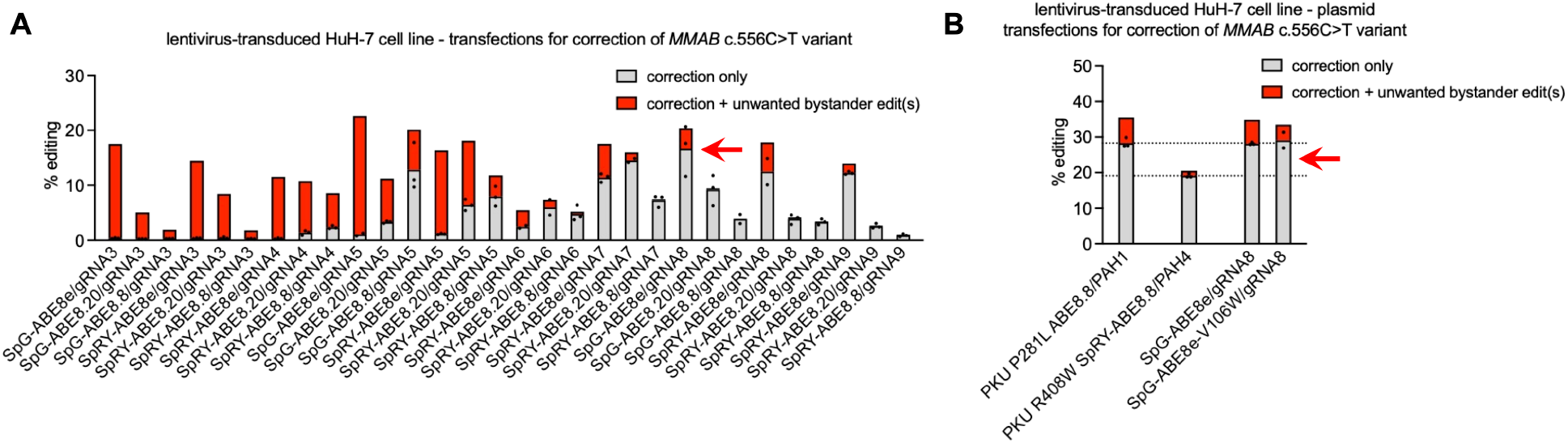
Optimization of adenine base editor/gRNA combinations for correction of the *MMAB* R186W variant. (A) Corrective editing (determined from genomic DNA) following transfection of lentivirus-transduced R186W HuH-7 cells with plasmids encoding various ABE/gRNA combinations (*n* = 3 biological replicates), calculated as the proportion of aligned sequencing reads with the indicated type of edits. “Correction only” refers to reads in which the R186W variant is corrected without base editing of one or more nearby adenines, i.e., bystander editing. SpG-ABE8e/gRNA8, indicated by the red arrow, performed the best out of all tested combinations. (B) Indicated by the red arrow, SpG-ABE8e-V106W (with the V106W variant in the deaminase domain, which eliminates gRNA-independent off-target editing and is more appropriate for clinical use) performed as well as the PKU P281L positive control that is known to produce efficient editing *in vivo* in mouse liver. The red horizontal dotted lines indicate the correction only editing levels for two validated, reference PKU variants P281L and R408W.

Because ABE8e has been reported to have gRNA-independent off-target RNA and DNA editing, the ABE8e-V106W variant, which largely eliminates the off-target editing of ABE8e,^16^ was used in a final plasmid transfection screen (**Figure 2B, Figure 3**). In the same transfection experiment, the validated solution^12^ for the *PAH* P281L variant (ABE8.8 with the published P281L-specific gRNA, designated PAH1) had similar corrective editing efficiency for the *PAH* P281L variant (**Figure 2B**, top horizontal dotted line) as compared to the corrective editing efficiency of SpG-ABEe-V106W/gRNA8 for the *MMAB* R186W variant.

**Figure 3.**
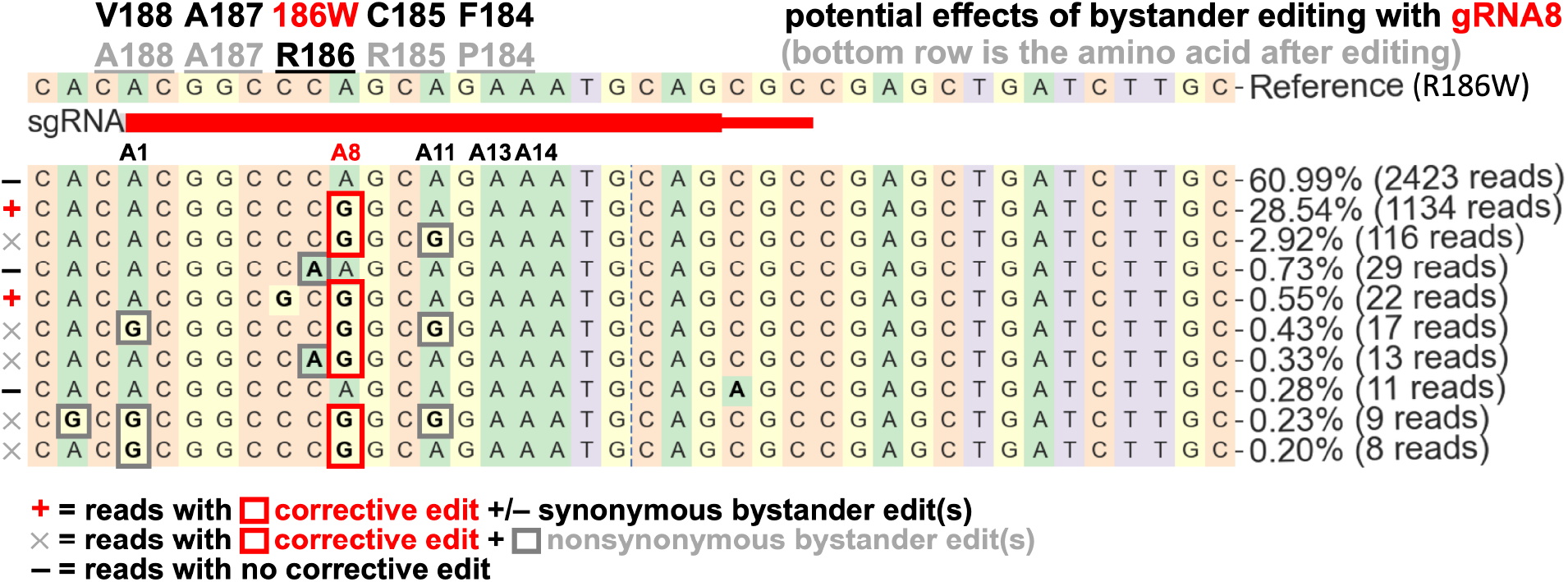
Corrective and bystander editing with SpG-ABE8e-V106W/gRNA8. Standard CRISPResso (http://crispresso2.pinellolab.org/submission) next-generation sequencing output showing the editing in a sample from the condition indicated by the red arrow in Figure 2B. The codons in the vicinity of the R186W site are indicated; the top-listed amino acid is the baseline/reference identity of the codon, and the bottom-listed amino acid is the one that results from base editing of the adenine in the codon. The red horizontal bar indicates the gRNA8 protospacer sequence, and the adjacent thin red box indicates the PAM sequence. All observed bystander editing by gRNA8 represents nonsynonymous changes that might affect the function of the MMAB protein.

We then tested gRNA8 in combination with NGC-ABE8e-V106W, an ABE with more specificity for the gRNA8 NGC PAM (see **Figure 1**),^4^ using *in vitro* transcribed mRNA encoding the ABE and chemically synthesized gRNA for mRNA/gRNA transfection (rather than plasmid transfection). We assessed two hybrid gRNA configurations of gRNA8 (“hyb16”, “hyb17”), with DNA nucleotide substitutions in the spacer sequence that have previously been reported to reduce both bystander editing and off-target editing by ABEs,^17^ along with the standard gRNA8 configuration (**Figure 4A**). The hyb16 configuration of gRNA8 has DNA nucleotide substitutions in positions 3, 4, and 5 of the spacer sequence, and the hyb17 configuration of gRNA8 has DNA nucleotide substitutions in positions 4, 5, and 6 of the spacer sequence. The hybrid gRNAs had substantially reduced bystander editing, resulting in an increased proportion of editing events with correction of the *MMAB* R186W variant only (the desired outcome) and surpassing the corrective editing efficiency by the validated solution for the *PAH* P281L variant.

**Figure 4.**
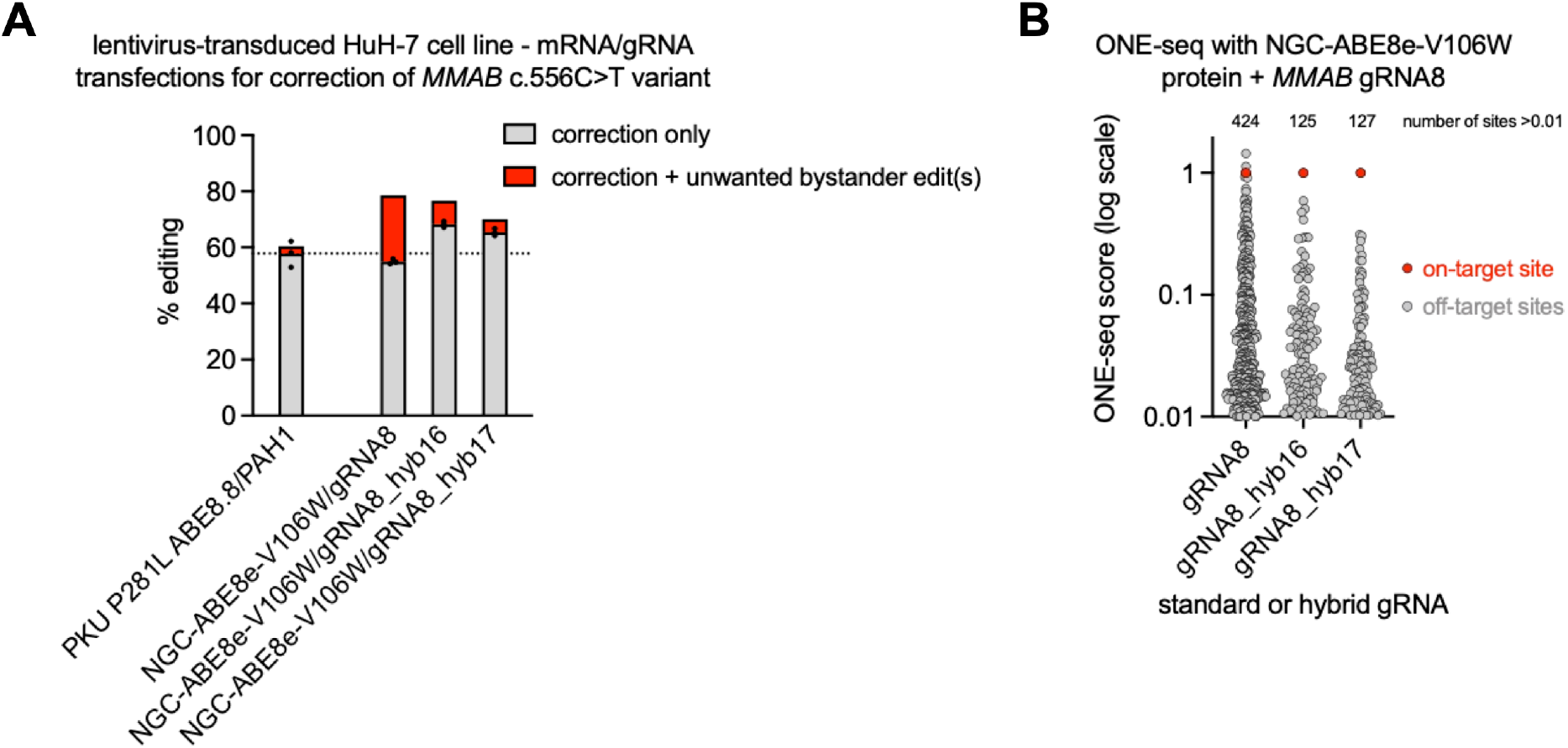
Further optimization of base editing with NGC-ABE8e-V106W/gRNA8. gRNA8 was tested in combination with NGC-ABE8e-V106W, an editor with more specificity for the gRNA8 NGC PAM, in lentivirus-transduced R186W HuH-7 cells. (A) Two hybrid guide RNA configurations of gRNA8 (hyb16, hyb17), with DNA nucleotide substitutions in the spacer sequence, were tested along with the standard gRNA8 configuration. NGC-ABE8e-V106W mRNA was co-transfected with chemically synthesized gRNA (*n* = 3 biological replicates). The hybrid gRNAs had substantially reduced bystander editing, resulting in an increased proportion of editing events with correction of the variant only. (B) ONE-seq assays showed the hybrid gRNAs to have less off-target potential than the standard gRNA, as assessed by the number of genomic sites with ONE-seq scores > 0.01.

To assess for off-target editing, we performed OligoNucleotide Enrichment and sequencing (ONE-seq), a homology-dependent biochemical assay that uses a synthetic human genomic library selected by sequence similarity to the protospacer/PAM sequence specified by the ABE/gRNA.^18,19^ We designed a library with sites in the reference human genome with up to five mismatches, or up to three mismatches plus up to two RNA or DNA bulges, to the on-target protospacer/PAM sequence. We performed the ONE-seq procedure with ribonucleoprotein complexes comprising recombinant NGC-ABE8e-V106W protein and the standard, hyb16, or hyb17 configuration of gRNA8. The ONE-seq assays generated rank-ordered lists of candidate (potential) off-target sites, with the on-target site set at a ONE-seq score of 1.0 and all other sites normalized to the on-target site (**Figure 4B**). Whereas standard gRNA8 had 424 candidate sites with ONE-seq scores > 0.01, the hybrid configurations of gRNA8 had 125–127 candidate sites with ONE-seq scores > 0.01, indicative of reduced potential for off-target editing. We used rhAmpSeq, a multiplex PCR method, to interrogate the top 119 ONE-seq-nominated candidate genomic sites in mRNA/gRNA-treated lentivirus-transduced R186W HuH-7 cells. Using individual targeted amplicon sequencing, we focused on the three sites with the most evidence of off-target editing with standard gRNA8, designated OT-11, OT-22, and OT-92; we also observed low-level bystander editing at the wild-type endogenous *MMAB* R186 site (which differs from the gRNA8 protospacer sequence by one nucleotide, the R186W variant adenine). OT-11 (located at chr7:113558333) is in an intergenic region far from any gene; OT-22 (chr16:48562359) is located in the *N4BP1* gene, which is involved in cytokine signaling; and OT-92 (chr1:17564859) is located in an intron of the *ARHGEF10L* gene, ≈20 kb away from exonic sequence in either direction. Both hybrid gRNAs greatly reduced editing at all four of these sites (**Figure 5**).

**Figure 5.**
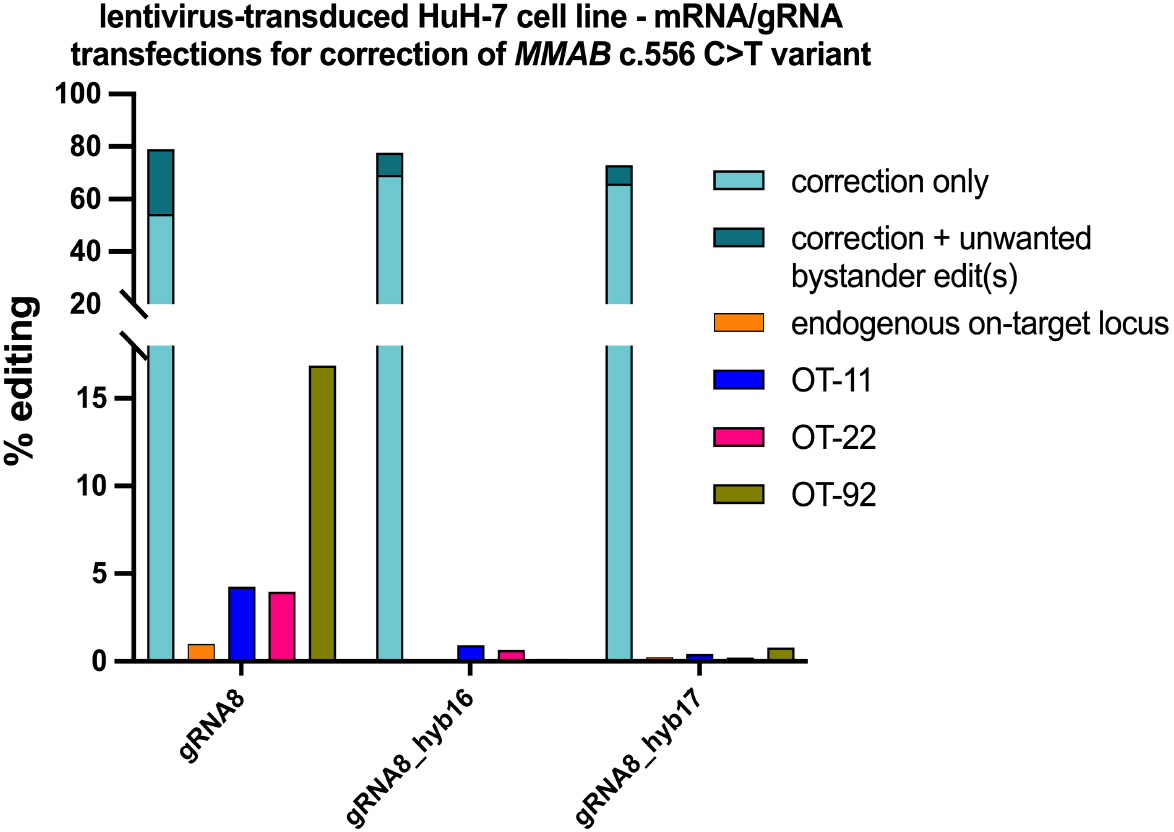
Reduction of off-target editing by hybrid gRNAs. (A) Two hybrid guide RNA configurations of gRNA8 (hyb16, hyb17) were tested along with the standard gRNA8 configuration in lentivirus-transduced R186W HuH-7 cells. NGC-ABE8e-V106W mRNA was co-transfected with chemically synthesized gRNA (*n* = 3 biological replicates). Individual targeted amplicon sequencing was performed at the on-target R186W site within the lentiviral cassette, the wild-type endogenous *MMAB* R186 site, and three verified sites of off-target editing.

Based on our on-target editing and off-target editing observations, we chose the hyb16 configuration of gRNA8 for further evaluation. We synthesized a version of the hyb16 gRNAs with a full set of chemical modifications that would be compatible with therapeutic use.^20^ We co-formulated this gRNA and NGC-ABE8e-V106W mRNA into LNPs with a well-established composition comprising the SM-102 ionizable lipid. We performed a dose-response experiment with these LNPs in lentivirus-transduced R186W HuH-7 cells (**Figure 6**). The LNPs achieved near-saturation editing at higher doses, with a calculated EC^50^ value ≈ 30 ng/mL, consistent with potencies observed with previous LNP test articles that demonstrated efficient *in vitro* and *in vivo* base editing.^4,12^

**Figure 6.**
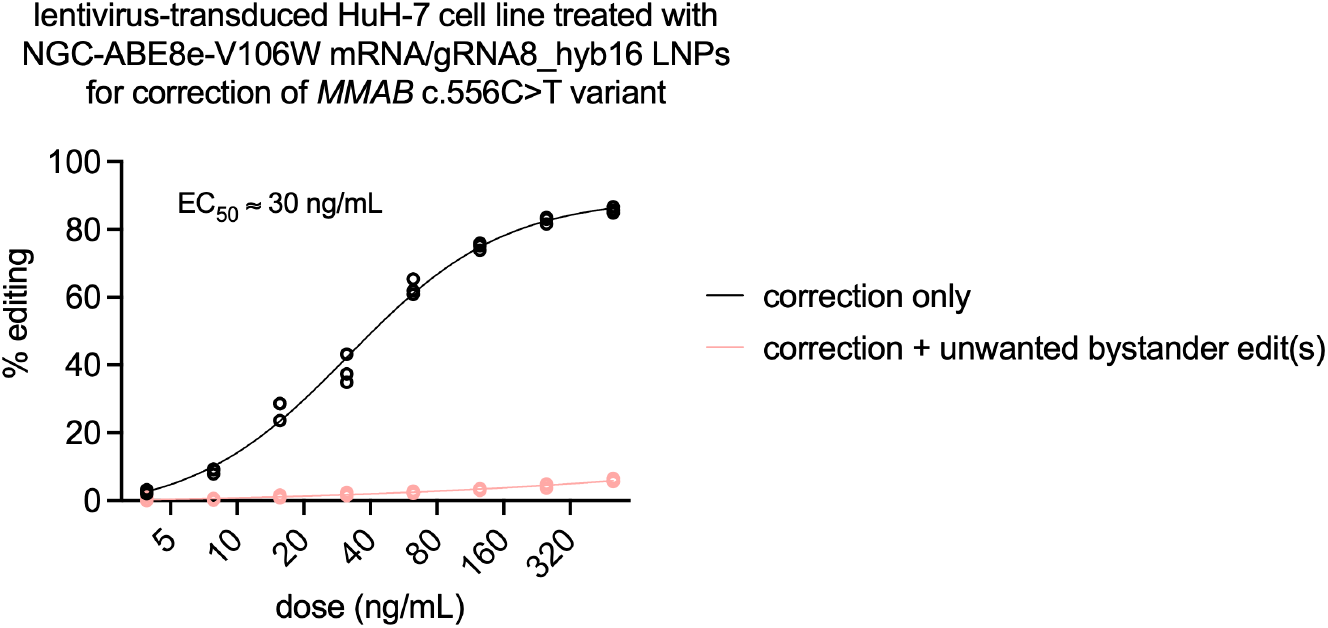
Dose-response study with LNPs. Corrective editing following transfection of lentivirus-transduced R186W HuH-7 cells with LNPs encapsulating NGC-ABE8e-V106W mRNA and the hyb16 configuration of gRNA8 (*n* = 3 biological replicates per dose). The wEC^50^ value was calculated from the best-fit curve.

## Discussion

*In vivo* gene editing, and particularly LNP-mediated adenine base editing, has taken enormous strides in the last few years, achieving clinical success in patients with familial hypercholesterolemia and atherosclerotic cardiovascular disease, alpha-1 antitrypsin deficiency, and CPS1 deficiency.^4,21,22^ The work described here supports the use of adenine base editing for the treatment of methylmalonic acidemia, cobalamin B type caused by the *MMAB* c.556C>T (R186W) variant. LNP-mediated delivery of an optimized mRNA/gRNA combination into the human liver could achieve the durable treatment of patients with MMAB disease caused by at least one copy of this variant. The potential for this therapy is enhanced by the reported positive selective advantage of corrected hepatocytes in an MMA mouse model following therapeutic gene insertion;^23^ if a similar phenomenon were to occur in the human liver, the therapeutic editing threshold for MMAB patients might be as low as a few percent, well within the reach of LNP-mediated adenine base editing.

Although the adenine base editing solution described here is directly relevant only to the *MMAB* c.556C>T (R186W) variant, a number of other reported pathogenic variants in *MMAB* as well as additional MMA genes are G>A or C>T substitutions that are potentially amenable to corrective adenine base editing. As we have previously reported, the U.S. Food and Drug Administration is amenable to “umbrella” clinical trials in which subjects with any of several closely related diseases (e.g., the urea cycle disorders) caused by any variant amenable to corrective adenine base editing could be enrolled in a single trial under a master protocol.^24^ The same consideration could apply to the MMAs and other organic acidemias, raising the prospect of accelerated approval of a platform of personalized editing therapies for these diseases. Thus, although the commercial potential for treatment of a single variant out of many in the MMAs, which are themselves rare diseases, might be limited, we anticipate that a general platform approach to correcting variants in MMAs and other IEMs will have substantial value in addressing the unmet medical need of affected patients.

## Methods

For base editing, a variety of adenine base editor (ABE)-expressing plasmids were used as previously described:^4^ ABE8.8-m (ABE8.8), SpG-ABE8.8, SpRY-ABE8.8, ABE8.20, SpG-ABE8.20, SpRY-ABE8.20, ABE8e, SpG-ABE8e, SpG-ABE8e-V106W, and SpRY-ABE8e. The pGuide plasmid (Addgene #64711) was used to express each accompanying gRNA (specific for the *MMAB* R186W variant) following subcloning of the oligonucleotide-synthesized gRNA sequence. Alternatively, *in vitro* transcribed mRNAs were produced and purified, as previously described: NGC-ABE8e-V106W, ABE8.8.^4,12^

Standard 100-mer gRNAs were chemically synthesized under solid phase synthesis conditions by a commercial supplier (Integrated DNA Technologies, or IDT) with end-modifications alone (standard configurations of gRNA3 to gRNA9, and hyb16 and hyb17 configurations of gRNA8), as well as a version of the hyb16 configuration of gRNA8 with heavy 2’-O-methylribosugar modification as previously described^17^: 5’-mA*mC*dG*dGdCCCAGCAGAAAUGCAGGUUUUAGAmGmCmUmAmGmAmAmAmUm AmGmCAAGUUAAAAUAAGGCUAGUCCGUUAUCAmAmCmUmUmGmAmAmAmAmA mGmUmGmGmCmAmCmCmGmAmGmUmCmGmGmUmGmCmU*mU*mU*mU-3’; where “m” and * indicate 2’-O-methylation and phosphorothioate linkage, respectively, and “d” indicates DNA nucleotide substitution. LNPs with the SM-102 ionizable lipid were formulated as previously described.^12^

HuH-7 human hepatoma cells were obtained from the Japanese Collection of Research Bioresources (JCRB) Cell Bank and maintained in culture with DMEM containing 1g/L glucose and supplemented with 10% FBS (Thermo Fisher). They were transduced with a lentiviral vector with a cassette harboring the *MMAB* R186W variant, the *PAH* (PKU) P281L variant, and the *PAH* (PKU) R408W variant, using a previously described strategy.^4^ The lentivirus-transduced HuH-7 cells were maintained until needed for transfection experiments. On the day prior to transfection, the HuH-7 cells were seeded on 24-well plates (Corning) at 1.2 × 10^5^ cells per well to achieve ≈80% confluence at the time of transfection. For plasmid transfections, at 3–4 hours after seeding, the cells were transfected at approximately 80–90% confluency with 1.5 μL of TransIT-LT1 transfection reagent (Mirus), 0.25 μg of base editor plasmid, and 0.25 μg of gRNA plasmid per well, according to the manufacturer’s instructions. For mRNA/gRNA transfections, in one tube, 3 µL of Lipofectamine MessengerMAX Transfection Reagent (Thermo Fisher Scientific) was added to 50 µL of Opti-MEM (Thermo Fisher Scientific) and incubated for 10 minutes; in another tube, 0.5 µg of mRNA and 0.5 µg of gRNA was suspended in a total of 50 µL of Opti-MEM. The diluted RNA was added to the tube of diluted MessengerMAX, and the subsequent RNA/MessengerMax/Opti-MEM mixture was incubated for 5 minutes at room temperature and then added dropwise onto the cells. For LNP transfection, LNPs were added at various doses (quantified by the total amount of RNA within the LNPs) directly to the media. For each type of transfection, cells were cultured for 72 hours after transfection, and then media were removed, cells were washed with 1× DPBS (Corning), and genomic DNA was isolated using the DNeasy Blood and Tissue Kit (Qiagen) according to the manufacturer’s instructions.

PCR amplification of individual target sequences (*MMAB* R186W or PKU P281L/R408W sequences in the lentiviral cassette) in genomic DNA samples from transfected lentivirus-transduced HuH-7 cells was performed using NEBnext High-Fidelity 2X PCR Master Mix (New England Biolabs) with locus-specific primers containing 5′ Nextera adaptor sequences (Illumina), followed by purification of the PCR amplicons with BioDynami Magnetic Beads for PCR purification (BioDynami) or the NGS Normalization 96-Well Kit (Norgen Biotek). A second round of PCR with the Nextera XT Index Kit V2 Set A and/or Nextera XT Index Kit V2 Set D (Illumina), followed by purification with the BioDynami Magnetic Beads or NGS Normalization 96-well kit, generated barcoded libraries, which were pooled and quantified using a Qubit 3.0 Fluorometer. After denaturation, dilution to 10 pM, and supplementation with 15% PhiX, the pooled libraries underwent next-generation sequencing on an Illumina MiSeq System.

The amplicon sequencing data were analyzed with CRISPResso2 (https://crispresso.pinellolab.partners.org/). rhAmpSeq multiplex PCR (IDT) with genomic DNA samples from transfected lentivirus-transduced HuH-7 cells was performed as previously described.^4^ ONE-seq was performed as previously described^4,12^ using recombinant NGC-ABE8e-V106W protein produced by GenScript.

## Acknowledgements

The authors gratefully acknowledge support from the NIH Common Fund Program, Somatic Cell Genome Editing, grants U19-NS132301 and U01-TR005355 (K.M., R.C.A.-N.), and a Cell and Gene Therapy Acceleration Grant from Children’s Hospital of Philadelphia (R.C.A.-N.).

